# Domain general mnemonic-attentional brain states fluctuate on sub-second timescales

**DOI:** 10.64898/2026.07.21.739572

**Authors:** Nicole M. Long

## Abstract

You remember what you attend to and you attend to what you remember. Memory is supported through engagement of mnemonic brain states, multivariate whole brain activity patterns that modulate down- stream processing and behavior. Attention can be focused externally to the environment or internally to thoughts and mental representations. To understand and promote successful cognition, it is critical to establish the extent to which the same brain states support both memory and attention processes. We recorded scalp EEG during three tasks in which we manipulated external and internal attention demands. We applied an independently-validated mnemonic state classifier to these data and find evidence in support of our hypothesis that memory encoding and retrieval states map onto the external/internal axis of attention. Furthermore, our findings reveal sub-second fluctuations in mnemonic states. These results demonstrate that domain general mnemonic states support attentional orienting and can be used to detect moment-to-moment shifts in attention.

## Introduction

You shouldn’t text and drive because of limited capacity attentional resources^1, 2^ – if your focus is on your phone, it isn’t on the road. However, in addition to the ability to focus your attention externally (to the buzz of a notification or the honk of a horn), you can also focus your attention internally, to mental representations, goals, and thoughts^3, 4^. To the extent that internal attention trades off with external attention, you also shouldn’t engage in mind-wandering while driving. The external/internal axis of attention may constitute a set of domain general, content-independent brain states that are linked with mnemonic brain states^5–7^. Memory encoding and memory retrieval are neurally distinct states that trade off^8, 9^, meaning that engaging in one state precludes engaging in the other, which has consequences for downstream processing and behavior^10–12^. Thus encoding/retrieval and external/internal attention may reflect common processes subject to the same limited capacity constraints. The aim of the present study is to elicit external and internal attention across a series of unique attentional tasks and to link the recruited neural substrates across attentional tasks and to mnemonic brain states. Determining the domain generality of attentional brain states and the extent to which these states overlap with mnemonic states is critical to understanding the role of both attention and memory across cognition.

External and internal attention may constitute domain general brain states. Brain states are temporally sustained whole brain activity/connectivity patterns that guide downstream processing and behavior^13–15^. Memory encoding and memory retrieval states recruit distinct hippocampal configurations^8^ and cortical activity patterns^10^ such that the two states trade off and cannot be engaged in simultaneously. Selective mnemonic state engagement impacts stimulus processing and subsequent memory^9–12^. External and internal attention may similarly constitute domain general brain states that support attentional orienting. External attention is the act of focusing on sensory information in the environment and is supported by the dorsal and ventral attention networks (DAN, VAN^16–18^) and spectral activity across multiple frequency bands (including theta [4-8Hz] and gamma [40Hz+])^19, 20^. Internal attention is the act of focusing on information stored within the mind and is primarily supported by the default mode network (DMN^21, 22^) and spectral activity in the alpha band (10-14Hz;^23–25^). Although task factors such as stimulus modality will modulate the downstream impacts of attention (e.g.^26^), content-independent attentional states should not vary across such factors^27^. Thus, although there is some evidence that external and internal attention can co-occur^28, 29^, external and internal attention may constitute distinct brain states.

Although the retrieval state (or mode,^30^) was initially proposed to be episodic-memory specific – a tonically maintained state that supports remembering an event situated within a specific time and place – increasing evidence suggests that the retrieval state extends beyond episodic memory and reflects internal attention^5, 7, 31–33^. The retrieval state is invoked in response to spatial attention^6^, semantic retrieval^34^ and working memory^35^ demands. The retrieval state recruits sustained engagement of a voltage topography consistent with engagement of the DMN^36^. Together, these findings suggest a strong overlap between mnemonic brain states and the external/internal axis of attention. Establishing the link between these brain states is critical to detecting, selecting, and correcting brain state engagement to support cognition.

Our hypothesis is that external and internal attention constitute neurally separable, domain general, content-independent brain states. The alternative hypothesis is that distinct external and internal processes are recruited depending on specific factors, including stimulus modality (e.g. visual vs. auditory), task structure (e.g. blocked vs. interleaved) and/or response demands. Our second hypothesis is that external and internal attention overlap with memory encoding and memory retrieval states, respectively. The alternative hypothesis is that mnemonic states are specifically recruited in response to episodic memory demands and thus are not engaged by attentional tasks lacking such demands. To test our hypothesis, we conducted a three task scalp electroencephalography (EEG) paradigm in which participants explicitly engaged external and internal attention while we varied task factors including stimulus modality, task structure, and response demands (Figure 1). We used a series of multivariate pattern analyses to assess external vs. internal attentional engagement and the degree to which these attentional demands modulated mnemonic state engagement.

**Figure 1.**
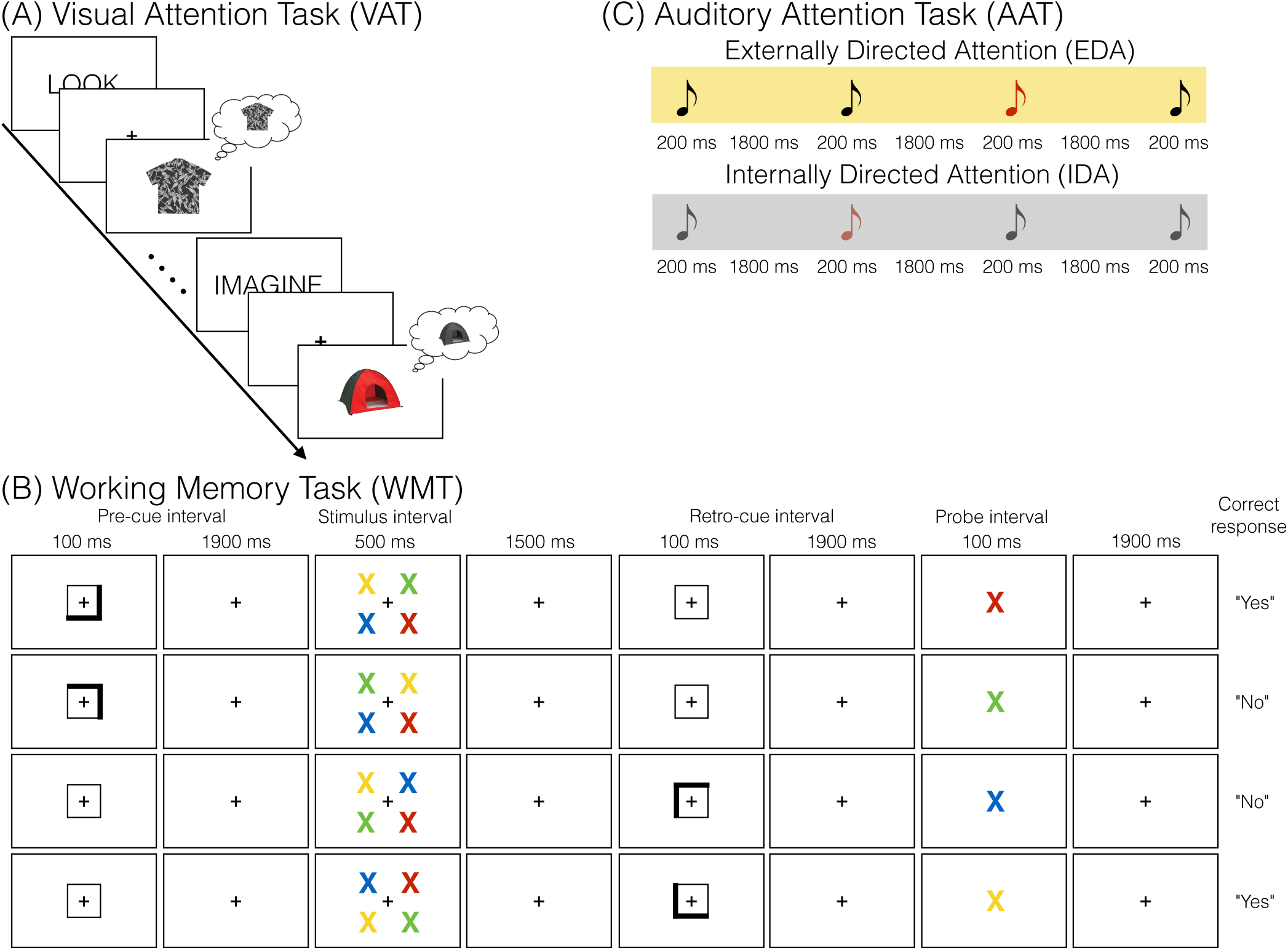
Task Design. **(A).** Visual Attention Task (VAT). Participants view color and grayscale images of common objects. Prior to each object participants are given an instruction cue to either “LOOK” at the presented object or “IMAGINE” the presented object in its opposite form – either in color or grayscale. In the example, participants should look at and focus on the greyscale shirt. Participants should imagine the colorful tent in grayscale. **(B)** Working Memory Task (WMT). Each trial begins with a 200 ms trial onset interval followed by an 1800 ms interstimulus interval (not shown). Participants were shown a directional cue during either the pre-cue interval or the retro-cue interval, but not both. Cue intervals are 100 ms in duration. Participants view four X’s, one of each: red, yellow, green, blue, in four locations during the 500 ms stimulus interval. Participants’ task is to respond as to whether the probe X matches (“yes”) or does not match (“no”) the pre/retro cued X. **(C)** Auditory Attention Task (AAT). Participants hear standard (black) and target (red) tones. All tones are played for 200 ms and separated by an 1800 ms inter-trial interval. In the externally directed attention (EDA) blocks (top), participants should attend to the tones and respond via keyboard when the target tone is played. In internally directed attention (IDA) blocks (bottom), participants should focus on their thoughts and not respond to the tones.

## Results

### Neurally dissociable external vs. internal attentional demands

Our first goal was to determine whether external and internal attention conditions are decodable within each task. We performed within-participant leave-one-run-out cross-validated-classification for each task. We trained each within-participant classifier to distinguish either (1) external (“LOOK”) vs. internal (“IMAGINE”) trials in the Visual Attention Task (VAT), (2) pre-cued vs. retro-cued trials in the Working Memory Task (WMT) or (3) externally directed attention (EDA) vs. internally directed attention (IDA) trials in the Auditory Attention Task (AAT). For each classifier, we averaged z-scored spectral power across the 0-2000 ms time interval of interest for that task. For the VAT and AAT, the interval of interest was the stimulus interval. For the AAT, we excluded target tone trials, thus auditory input and response demands were identical across the two conditions. For the WMT, the interval of interest was the pre-cue interval on pre-cued trials and the retro-cued interval on retro-cued trials, meaning that the visual input was identical across the two conditions. The average classification accuracy for all three tasks was significantly above chance as determined via permutation procedure (Table 1, Figure 2A-C, Top Row).

**Figure 2.**
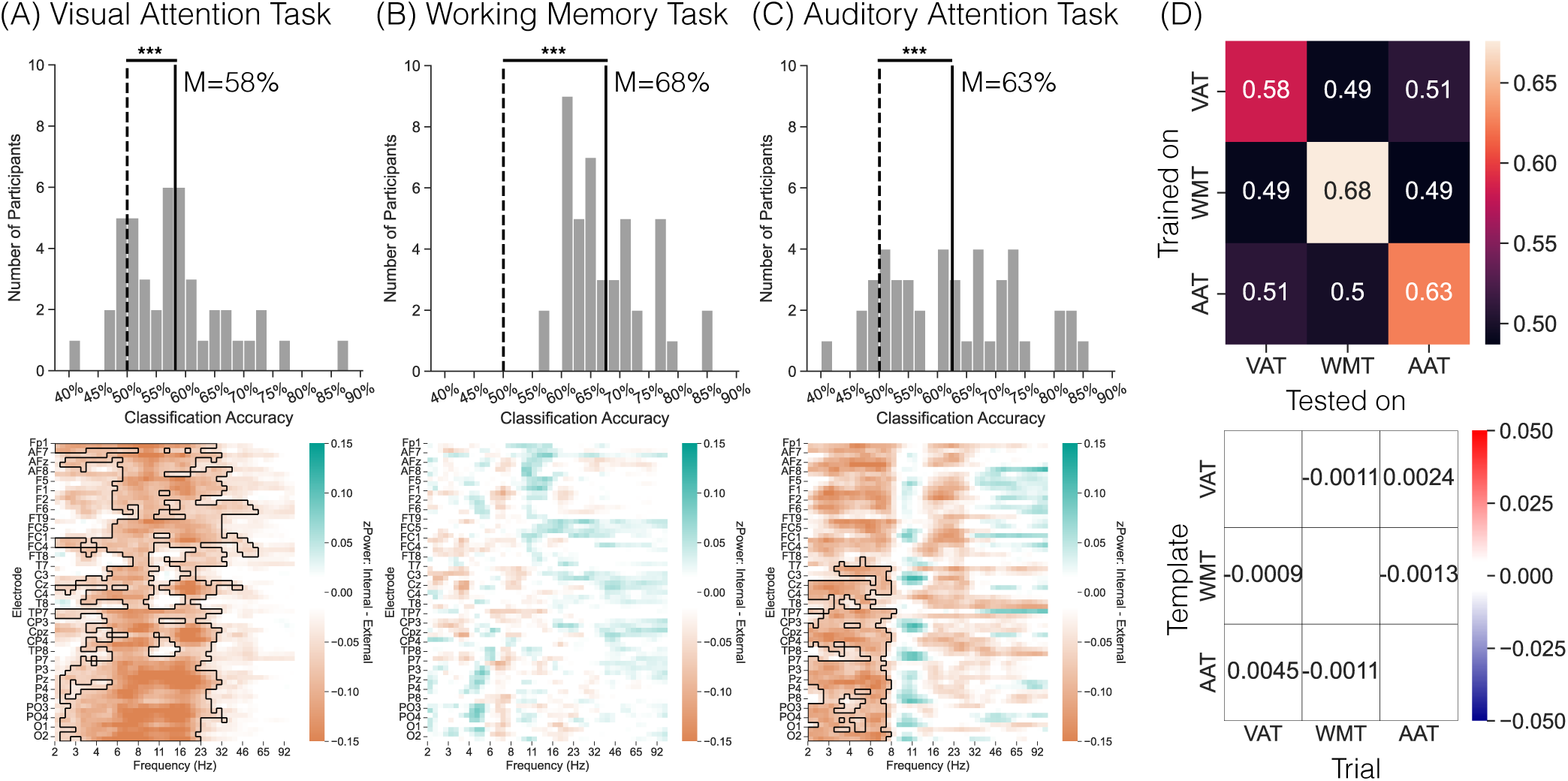
Within- and cross-task attention comparisons. We averaged spectral signals across the 2000ms stimulus (Visual Attention Task, VAT; Auditory Attention Task, AAT) or pre/retro-cue interval (Working Memory Task, WMT). **(A-C)** Top Row: We conducted leave-one-run-out cross-validated classification on external and internal trials for each of the three tasks. We trained classifiers to distinguish LOOK (external) vs. IMAGINE (internal) VAT trials, pre-cued (external) vs. retro-cued (internal) WMT trials, or externally directed attention (EDA) vs. internally directed attention (IDA) AAT trials. Target tone trials were excluded. Solid vertical bars indicate mean classification accuracy. Dashed vertical bars indicate chance classification accuracy as determined by permutation procedures. Classification accuracy was significantly above chance in each of the three tasks. *** *p <* 0.0001. Bottom Row: electrode *×* frequency spectrograms of the univariate contrast of the conditions upon which the classifier was trained and tested. Teal indicates greater zPower for the internal (imagine, retro-cued, IDA) condition relative to the external (look, pre-cued, EDA) condition. Orange indicates greater zPower for the external relative to the internal condition. Boxes indicate significant clusters following multiple comparisons correction. **(D)** Top Row: We trained within-participant classifiers to distinguish external/internal conditions in one task (y-axis) and tested each classifier to distinguish the analogous conditions in another task (x-axis). Colormap indicates classification accuracy. Values along the diagonal reflect within-task classification accuracy and are identical to the means shown in the A-C histograms. Bottom Row: We correlated trial-averaged template spectral patterns in one task (y-axis) with trial-level spectral patterns in another task (x-axis). Colormap indicates zRho values. Same-task comparisons (diagonal) were not performed.

**Table 1.** Within-task attention state classification accuracy.

| Task | M | SD | $t_{43}$ | $p$ | $d$ | N |
| --- | --- | --- | --- | --- | --- | --- |
| VAT | **58.25%** | **8.930%** | **6.004** | **<0.0001** | **1.305** | 26 |
| WMT | **67.61%** | **6.650%** | **17.22** | **<0.0001** | **3.730** | 44 |
| AAT | **62.53%** | **11.14%** | **7.443** | **<0.0001** | **1.585** | 27 |

*Bold values indicate $p < 0.05$ . N indicates the number of participants with above-chance accuracy*

To directly compare decoding performance across the three tasks, we conducted a 1*×*3 rmANOVA. We find a significant effect of task (*F* _2,86_=11.31, *p<*0.0001, *η_p_*^2^= 0.2083) driven by significantly greater classification accuracy on the WMT (M=67.61%, SD=6.730%) compared to the VAT (M=58.25%, SD=9.030%; *t* _43_=5.363, *p<*0.0001, *d* =0.8084, BF=5901) and the AAT (M=62.53%, SD=11.28%; *t* _43_=2.566, *p*=0.0120, *d* =0.3820, BF=2.787). Classification accuracy did not significantly differ between the VAT and AAT (*t* _43_=2.000, *p*=0.0519, *d* =0.3015, BF=0.9993).

Together, these findings indicate that it is possible to distinguish externally vs. internally directed attention in visual or auditory attention tasks, when tasks are blocked or interleaved, and in the presence or absence of response demands. That we found exceptionally robust task decoding in the WMT is notable given that the visual input is matched between the two attention conditions. This finding suggests that participants engage distinct neural substrates to support preparatory allocation of attention during the pre-cue interval vs. to update the contents of working memory during the retro-cue interval.

### Distinct univariate signals underlie each attention task

Having demonstrated that external and internal attention demands are distinguishable within each task, we next sought to generalization of these signals across tasks. To the extent that participants recruit a single domain general external (or internal) attentional state, we should find robust cross-task decoding. Alternatively, to the extent that participants recruit task-specific external (or internal) attention processes, cross-task decoding may fail. For instance, participants may recruit different mechanisms to support mental imagery of grayscale object images in the VAT vs. focusing on their thoughts in the IDA blocks of the AAT. In such a case, the features that distinguish external from internal attention would differ across tasks, preventing cross-task generalization.

We conducted within-participant, cross-task classification for the six pairs of tasks. We averaged classification accuracy values across pairs of the same tasks (e.g. trained on VAT, tested on WMT and trained on WMT and tested on VAT). We find that classification accuracy does not significantly differ from chance for any of the pairs of tasks (as determined by permutation procedure; Table 2, Figure 2D).

**Table 2.** Cross-task attention state classification accuracy and template correlations.

| Task | | M | SD | $t_{43}$ | $p$ | $d$ | BF |
| --- | --- | --- | --- | --- | --- | --- | --- |
| VAT-WMT | Classification Accuracy | 48.95% | 4.500% | 1.613 | 0.1141 | 0.2459 | 0.5397 |
|  | Template Correlation | 0.0045 | 0.0236 | 1.270 | 0.2109 | 0.1907 | 0.3455 |
| WMT-AAT | Classification Accuracy | 49.24% | 7.220% | 0.6869 | 0.4958 | 0.1048 | 0.2038 |
|  | Template Correlation | -0.0004 | 0.0182 | 0.1428 | 0.8871 | 0.0220 | 0.1648 |
| AAT-VAT | Classification Accuracy | 50.88% | 4.480% | 1.433 | 0.1592 | 0.2158 | 0.4219 |
|  | Template Correlation | 0.0013 | 0.0217 | 0.3913 | 0.6975 | 0.0599 | 0.1755 |

Given the robust within-task decoding, the failure of cross-task decoding implies that non-overlapping neural correlates are recruited for each of the three tasks. To examine the neural correlates that underlie each task contrast, we conducted an exploratory univariate analysis in which we compared z-scored power (zPower) for external (LOOK, pre-cued, EDA) vs. internal (IMAGINE, retro-cued, IDA) trials in each task (Figure 2A-C, Bottom Row). In the VAT, we find significantly greater low frequency (2-23Hz) zPower broadly across the scalp for external relative to internal trials. In the WMT, no clusters survived multiple comparisons correction. Finally, in the AAT, we find significantly greater posterior theta (2-8Hz) zPower for external relative to internal trials.

We directly compared spectral patterns across the pairs of tasks. We compared trial level spectral activity in one task to averaged or “template” external and internal patterns in another task using Pearson correlations (see Methods). None of the comparisons yielded zRho values that significantly differed from zero (Table 2). Thus, the spectral signals that support external vs. internal processing differ across tasks.

### Mnemonic brain states are selectively recruited for all attention tasks

Our central goal was to apply the mnemonic state classifier to the three tasks in the current study. Across all tasks, we expected to find increased encoding state evidence during external trials and increased retrieval state evidence during the internal trials. In all analyses, mnemonic state evidence served as the dependent variable. Due to the structure of our classifier, positive evidence values indicate greater evidence for the encoding state whereas negative values indicate greater evidence for the retrieval state.

We tested the cross-study mnemonic state classifier on the stimulus interval (0-2000 ms) of the VAT data separately for each instruction (LOOK/external, IMAGINE/internal; Figure 3A). We conducted a 2 (instruction) *×* 20 (twenty 100 ms time intervals) rmANOVA. We find a significant main effect of instruction (*F* _1,43_=5.133, *p*=0.0286, *η_p_*^2^=0.1066). Mnemonic evidence is significantly more positive for LOOK (M=-0.0002, SD=0.0288) relative to IMAGINE (M=-0.0204, SD=0.0372) trials. We find a significant main effect of time interval (*F* _19,817_=57.35, *p<*0.0001, *η_p_*^2^=0.5715). We find a significant instruction by time interaction (*F* _19,817_=2.423, *p*=0.0006, *η_p_*^2^=0.0533) whereby mnemonic state evidence is significantly more positive for LOOK (M=-0.0288, SD=0.0466) relative to IMAGINE (M =-0.0602, SD=0.0620) trials in the second half of the stimulus interval (1000-2000 ms; *t* _43_=2.446, *p*=0.0186, *d* =0.3688, BF=2.326), but not the first half (LOOK, M=0.0284, SD=0.0330, IMAGINE, M=0.0194, SD=0.0391; 0-1000 ms; *t* _43_=1.286, *p*=0.2053, *d* =0.1939, BF=0.3520). Thus, although both tasks recruit the encoding state (positive mnemonic state evidence) early in the interval and the retrieval state (negative mnemonic state evidence) late in the interval, the instruction to imagine an object stimulus recruits the retrieval state to a greater degree than the instruction to look at an object stimulus.

**Figure 3.**
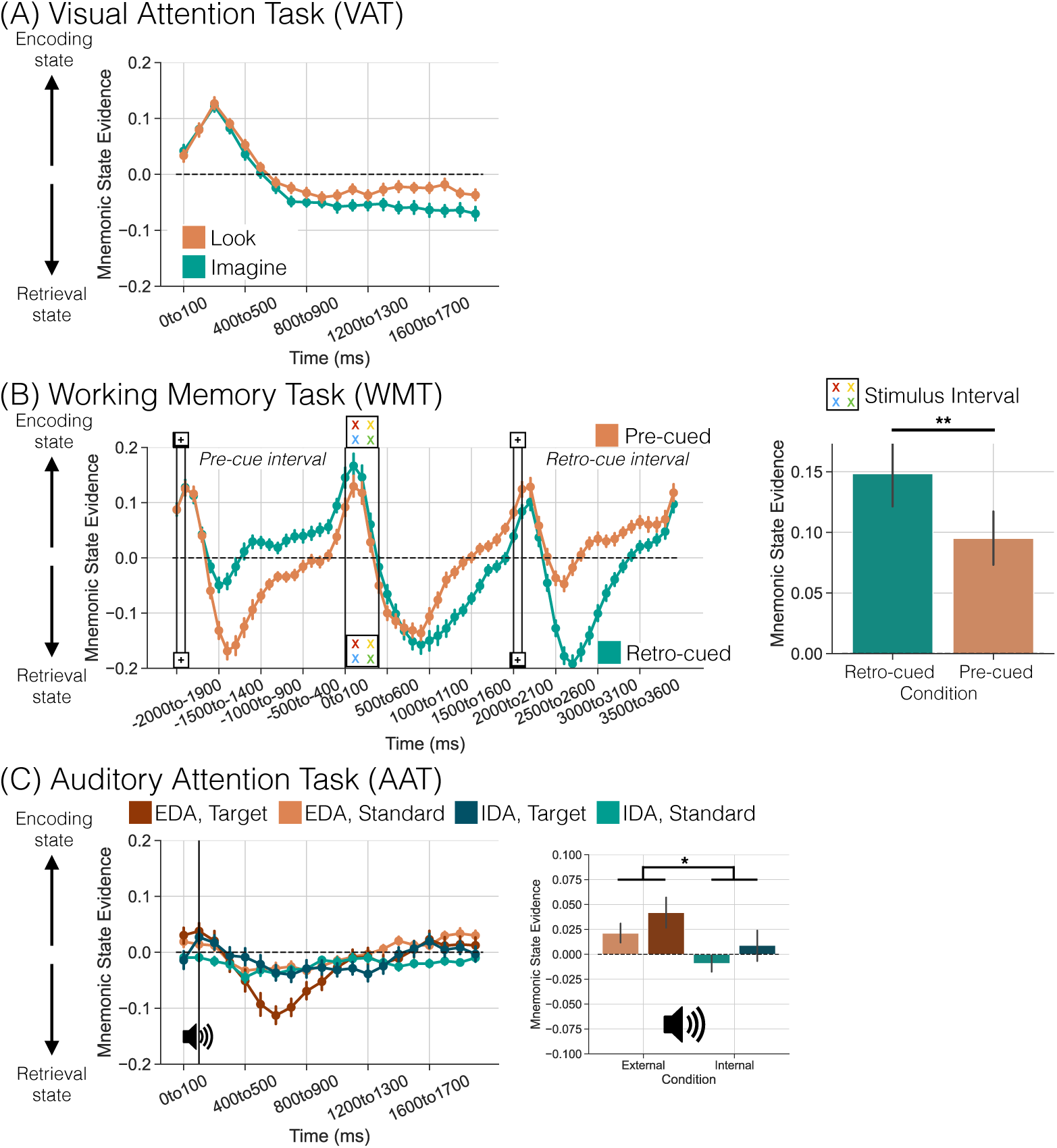
Mnemonic state engagement across three attentional tasks. Each figure shows mnemonic state evidence over time as a function of internal (teal) and external (orange) conditions in each task. Positive values reflect greater evidence for the encoding state whereas negative values reflect greater evidence for the retrieval state. **(A)** Mnemonic state evidence is greater for the external “LOOK” trials relative to the internal “IMAGINE” trials during the second half of the stimulus interval in the VAT. **(B)** Left: Mnemonic state evidence across the pre-cue, stimulus, and retro-cue intervals in the WMT. Pre-cued trial structure is illustrated on the top of the graph and represented by the orange line. Retro-cued trial structure is illustrated on the bottom of the graph and represented by the teal line. Mnemonic state evidence on retro-cued trials is greater during the pre-cue interval (−2000 to 0ms) relative to the retro-cue interval (2000 to 4000ms). Vertical lines indicate onset and offset of cue and stimulus intervals. Right: Mnemonic state evidence during the stimulus interval (0 to 500ms) is greater during retro-cued trials relative to pre-cued trials. **(C)** Left: Mnemonic state evidence across the stimulus (200 ms) and inter-trial interval (200 to 2000ms) as a function of block (external, orange, EDA; internal, teal, IDA) and tone type (target, darker hues; standard, lighter hues). Mnemonic state evidence varies over time as a function of block and tone type. Right: Mnemonic state evidence during the stimulus interval (0 to 200ms) is greater during EDA (orange) compared to IDA (teal) trials. Error bars represent standard error of the mean. * *p <* 0.05; ** *p <* 0.01

We next tested the cross-study classifier on the stimulus and cue intervals in the WMT. First, we assessed classification evidence during the stimulus (0-500 ms) interval separately as a function of cue type (precued vs. retro-cued; Figure 3B, right panel). To the extent that participants selectively attend to one of the four display items on pre-cued trials, the effective set size on those trials should be one rather than four. We have previously found greater recruitment of the encoding state as set size increases^35^. Therefore, we should find more encoding state evidence on retro- relative to pre-cued trials. We conducted a 2 (cue type) *×* 5 (five 100 ms time intervals during the stimulus interval) rmANOVA. We find a significant main effect of cue type (*F* _1,43_=8.139, *p*=0.0066, *η_p_*^2^=0.1591). Mnemonic state evidence is significantly more positive for retro-cued (M=0.1004, SD=0.1264) relative to pre-cued (M=0.0634,SD=0.1040) trials. We find a significant main effect of time interval (*F* _4,172_=88.00, *p<*0.0001, *η_p_*^2^=0.6717). We do not find a significant instruction by time interaction (*F* _4,172_=1.509, *p*=0.2017, *η_p_*^2^=0.0339). These results suggest that although the number of visually presented stimuli is held constant at four items, the pre-cue functionally decreases the set size from four to one such that there is greater encoding state engagement (positive mnemonic state evidence) for retro-cued relative to pre-cued trials.

Second, we compared mnemonic state evidence across the pre-cue and retro-cue intervals exclusively for retro-cued trials (Figure 3B, left panel, teal line). The logic of this analysis is that there should be little attention deployment during the pre-cue interval on retro-cued trials as there is no cue during that interval. In contrast, attention should be directed to the internally maintained array during the retro-cue interval. Therefore, we should find more retrieval state evidence (negative mnemonic state evidence) during the retro-cue interval relative to the pre-cue interval. We conducted a 2 (interval) *×* 20 (twenty 100 ms time intervals) rmANOVA. We find a significant main effect of interval (*F* _1,43_=62.09, *p<*0.0001, *η_p_*^2^=0.5908). Mnemonic state evidence is significantly more negative for retro-cue (M=-0.0293, SD=0.0387) relative to pre-cue (M=0.0356, SD=0.0363) intervals. Follow-up one sample *t* -tests vs. zero revealed that mnemonic state evidence is significantly positive for the pre-cue interval (*t* _43_=6.514, *p<*0.0001, *d* =0.9807, BF=2.169*×*10^5^) and significantly negative for the retro-cue interval (*t* _43_=5.034, *p<*0.0001, *d* =0.7571, BF=2159). We find a significant main effect of time interval (*F* _19,817_=45.70, *p<*0.0001, *η_p_*^2^=0.5311). We find a significant interval by time interaction (*F* _19,817_=23.19, *p<*0.0001, *η_p_*^2^=0.3504) driven by the largest dissociations in the middle (500-1500 ms) of the interval, encompassing the ISI between cue and stimulus/probe. Thus, in the absence of specific spatial cues (pre-cue interval), participants engage the encoding state, likely to prepare for the upcoming stimulus display. Upon receiving a cue (retro-cue interval), however, participants engage the retrieval state, likely to update their working memory storage and attend selectively to the cued item.

Finally, we tested the cross-study classifier on the stimulus interval (0-2000 ms) of the AAT separately for each task block (EDA, IDA) and tone condition (standard, target; Figure 3C, left panel). Insofar as external attention is recruited in response to both top-down demands to focus on the tones and bottom-up stimulus features, such as salience of the target tone, we should find differential mnemonic state recruitment as a function of both factors. Specifically, we should find greater encoding state recruitment during the EDA vs. the IDA task. We should additionally find greater encoding state recruitment during target vs. standard trials. We conducted a 2 (task) *×* 2 (tone) *×* 20 (twenty 100 ms time intervals) rmANOVA. We did not find significant main effects of task or tone as we anticipated (Table 3). Instead, we found that all interactions were significant, prompting a series of follow-up analyses.

**Table 3.** Impact of task, tone, and time on mnemonic state evidence in the AAT.

| Effect | df | $F$ | $p$ | $\eta_p^2$ |
| --- | --- | --- | --- | --- |
| Main effect of Task | 1,43 | 0.3625 | 0.5503 | 0.0084 |
| Main effect of Tone | 1,43 | 0.9606 | 0.3325 | 0.0219 |
| Main effect of Time | 19,817 | **14.76** | **< 0.0001** | **0.2555** |
| Interaction of Task × Tone | 1,43 | **7.950** | **0.0072** | **0.1560** |
| Interaction of Task × Time | 19,817 | **4.591** | **< 0.0001** | **0.0965** |
| Interaction of Tone × Time | 19,817 | **3.404** | **< 0.0001** | **0.0734** |
| Interaction of Task × Tone × Time | 19,817 | **2.276** | **0.0015** | **0.0503** |

*Bold values indicate $p < 0.05$*

We first assessed mnemonic state evidence over time exclusively for standard tone trials (Figure 3C, left panel, lighter lines). This approach leverages the same data utilized for the within-participant classification analysis in which target tones were excluded. We conducted a 2 (task) *×* 20 (twenty 100 ms time intervals) rmANOVA. We find a significant main effect of task (*F* _1,43_=7.195, *p*=0.0103, *η_p_*^2^=0.1433) driven by significantly more positive mnemonic state evidence for the EDA (M=-0.0014, SD=0.0234) compared to the IDA (M=-0.0206, SD=0.0269) task blocks. We find a significant main effect of time interval (*F* _19,817_=7.149, *p<*0.0001, *η_p_*^2^=0.1426). We find a significant interaction between task and time interval (*F* _19,817_=3.784, *p<* 0.0001, *η_p_*^2^=0.0809), driven by task dissociations early (0-300 ms) and late (1300-2000 ms) in the stimulus interval. Early and late in the stimulus interval, there is relatively more encoding state evidence for EDA blocks and relatively more retrieval state evidence for IDA blocks.

We next assessed mnemonic state evidence over time exclusively for target tone trials (Figure 3C, left panel, darker lines). We conducted a 2 (task) *×* 20 (twenty 100 ms time intervals) rmANOVA. We did not find a significant main effect of task (*F* _1,43_=0.6251, *p*=0.4335, *η_p_*^2^=0.0143). We find a significant main effect of time interval (*F* _19,817_=10.37, *p<*0.0001, *η_p_*^2^= 0.1943). We find a significant interaction between task and time interval (*F* _19,817_=3.416, *p<*0.0001, *η_p_*^2^=0.0736), driven by more negative mnemonic state evidence for EDA (M=-0.0208, SD=0.0533) compared to IDA (M=-0.0115, SD=0.0421) task blocks around 500-700ms following stimulus onset. We would have anticipated more positive mnemonic state evidence for the EDA blocks, insofar as participants attend externally and especially to the oddball target tone. That we find the opposite for the middle interval within the trial suggests that participants may rapidly modulate mnemonic state engagement depending on task demands and stimulus properties.

We had anticipated *a priori* that the bottom-up effects of the tone may be time limited as tones are only presented for 200 ms. Specifically, external attention to the target tones may only persist for a few hundred milliseconds, whereas top-down driven external attention in response to the task block instructions should be sustained throughout the stimulus interval. Therefore, we performed a targeted investigation of mnemonic state evidence, specifically on z-scored power averaged across the 0-200 ms following stimulus onset (Figure 3C, right panel). We conducted a 2*×*2 rmANOVA with task block and tone condition as factors. We find a significant main effect of task (*F* _1,43_=5.935, *p*=0.0191, *η_p_*^2^=0.1213) driven by more positive mnemonic state evidence on EDA (M=0.0314, SD=0.0674) compared to IDA (M=-0.0001, SD=0.0570) trials. Neither the main effect of tone (*F* _1,43_=1.934, *p*=0.1714, *η_p_*^2^=0.0430) nor the interaction of task and tone (*F* _1,43_=0.0161, *p*=0.8995, *η_p_*^2^=0.0004) were significant. These results suggest that top-down influences may exert a stronger influence on mnemonic state engagement relative to bottom-up stimulus salience.

Together, our cross-study mnemonic state analyses demonstrate that internal attention demands recruit the retrieval state whereas external attention demands recruit the encoding state, across different experimental paradigms and independent from stimulus modality, task structure, and response demands.

## Discussion

The objective of this study was to test the hypotheses that external and internal attention constitute neurally separable, domain general, content-independent brain states which overlap with memory encoding and memory retrieval states, respectively. We conducted three tasks that modulate external vs. internal attention while recording scalp electroencephalography (EEG). Task factors including stimulus modality, task structure, and response demands varied across the tasks. We find robust within-task attention state decoding meaning that spectral patterns across the scalp are consistently distinct between external and internal manipulations; however, we failed to find cross-task attention state generalization meaning that the underlying patterns that differentiate external vs. internal attention conditions differ as a function of task factors. Critically, we find that our putative episodic mnemonic state classifier is consistently modulated by external vs. internal attentional demands across all tasks. Together, these results indicate that mnemonic states index domain general attentional states, bridging across task factors. Mnemonic state evidence can thus be used to detect how attention is oriented on a moment to moment basis.

We show that external and internal attention demands are dissociable within visual, working memory, and auditory tasks and selectively engage mnemonic brain states. Multivariate machine learning classifiers can decode whether participants are asked to look at or imagine visual object stimuli presented in color or grayscale (visual attention task, VAT), perform a pre- vs. retro-cued working memory task (WMT), or attend to or ignore auditory tones (auditory attention task, AAT). As perceptual input and motor output were matched across conditions within each task, low-level information is unlikely to have enabled the classifiers to distinguish the attention conditions. However, within-task decoding likely reflects task-specific information as we failed to find any evidence that external and internal signals generalized across tasks. In contrast, we found strong support for our central hypothesis that mnemonic brain states reflect the external/internal axis of attention^3^. Both the memory encoding and memory retrieval state were robustly engaged across all tasks despite the variable task factors and the fact that none of the tasks require longterm episodic memory retrieval, originally proposed to selectively engage the retrieval mode^30^.

External vs. internal attention to visual stimuli are dissociable and differentially recruit mnemonic brain states. We designed the VAT to mirror our mnemonic state task in terms of stimuli and trial timing (e.g.^12^). We expected the “LOOK” cue to recruit external attention whereas we expected the “IMAGINE” cue to recruit internal attention. The spatially broad low frequency activity increases for external relative to internal trials is partly consistent with this expectation given evidence that signals within the theta band (4-8Hz) track attentional selection of perceptual inputs^37–39^ and that we find significantly more positive mnemonic state evidence (evidence for encoding) for external relative to internal trials late in the stimulus interval. However, the limited external condition increases in the alpha band (10-14Hz) are inconsistent with evidence that alpha power increases during imagery and internal tasks relative to perception and external tasks^23, 40, 41^. Corroborating the univariate effects, we find that although the two conditions are dissociable, both engage the retrieval state. We speculate that these results reflect how participants approached the task and the relative attentional demands. Of the 26 participants who reported using any strategy during the VAT, around 20% (n=6) reported internally vocalizing to themselves during both conditions. We expect such a process to recruit the retrieval state given that this brain state supports WM maintenance^35^ which encompasses verbal rehearsal^42^. Participants may subvocalize the item category regardless of instruction, recruiting internal attention even in the putative external condition. Additionally, to the extent that attention is recruited in response to competition^43^, there may be insufficient competition in the VAT to necessitate strong selection processes. Selective attention recruits alpha power^44, 45^, with greater recruitment as distractor strength increases^46^. The lack of distractors in the present task may limit the amount of attention required and thus effects within the alpha band. We expect that with more perceptual load or external attention demands^23, 47, 48^, for instance, use of more complex visual stimuli, reliance on external attention should increase with a concomitant increase in encoding state evidence and decrease in alpha power.

Mnemonic states are robustly modulated by each trial segment in the WMT. In the original fMRI WMT study^49^, both overlapping and distinct regions supported external vs. internal attention. Given the limited spatial resolution of scalp EEG, we cannot replicate region-level effects; however, by leveraging scalp EEG’s high temporal resolution, we were able to target specific segments within each trial and revealed sub-second mnemonic state fluctuations. We found highly robust within-participant decoding of the precue interval on pre-cued trials (putative external) vs. the retro-cue interval on retro-cued trials (putative internal). However, the group-level spectrogram of the external vs. internal WMT contrast revealed no significant clusters, indicating a high degree of cross-participant variability. The majority of the signal in the two conditions is likely driven by shared processes of reflexive cueing, attentional orienting, selecting, and maintenance^2^ which, when combined with variable strategy use, may account for the lack of group-level univariate dissociations. Participants reported naming colors (n=19, 50%), chunking the display stimuli (n=7, 18%) or some other strategy (n=13, 33%). Mnemonic state evidence is similarly decreased across the same two conditions, suggesting that both recruit internal attention. Although these findings contradict the notion that pre-cues recruit external attention, they are consistent evidence showing retrieval state engagement during the delay interval of a Posner spatial cueing task^6^. We speculate that participants recruit internal attention to maintain or selectively attend to the cued location for both pre-cued and retro-cued trials. The robust dissociation in mnemonic state engagement for the two intervals on retro-cued trials supports this interpretation. Specifically, we find greater retrieval state evidence during the retro-cue interval of retro-cued trials relative to the pre-cue interval of retro-cued trials. Thus, with the high temporal resolution of scalp EEG, we have revealed that both putative external and internal attention can be recruited in a matter of seconds within the same trial. Prior fMRI evidence for co-occurrence of external and internal attention may therefore reflect the blurring of temporally distinct signals. Given that we find greater retrieval state evidence for retro-cued vs. pre-cued trials within the retro cue interval, condition dissociations are not solely driven by which interval is under consideration. Our interpretation is that retrieval state evidence during the retro-cue interval on retro-cued trials reflects a “focus of internal attention highlighting one of several representations”^50^. Namely, on retro-cued trials, participants begin the interval by maintaining four stimuli and then update what is maintained by removing three of the four stimuli based on the retro-cue. Taken together, these findings show that the mnemonic state classifier tracks moment to moment changes in attentional states.

External vs. internal attention to auditory stimuli are dissociable and differentially recruit mnemonic brain states. We selected the AAT from prior investigations of external/internal attention^24, 25^. Kam and colleagues (2018) showed that theta power in the AAT increases for externally directed attention (EDA) and alpha power increases for internally directed attention (IDA) which we largely replicate here. We find theta power increases and encoding state engagement for EDA trials. Although not significant after multiple comparisons correction, we find numeric increases in alpha power for IDA trials, along with significant retrieval state engagement. However, we find unexpected temporally dissociable mnemonic state modulations as a combined function of top-down task goals to attend and bottom-up stimulus salience. Our *a priori* expectation was that target or oddball tones would recruit the encoding state to a greater degree than standard tones, given that highly salient stimuli should capture involuntary attention and associated attentional networks^16, 18^. Instead, we found greater *retrieval* evidence *following* target tones during EDA trials. We speculate that participants are able to modulate mnemonic state engagement on sub-second timescales in response to task structure. In the EDA blocks, participants presumably begin the trial in a high vigilance state in anticipation of the target tone. However, once the target tone is played, participants are likely aware that there will be several hundred milliseconds before any tone, and potentially more than several seconds before a target tone, will next be played. Thus, participants may disengage external attention and potentially even engage internal attention following the offset of the tone. It is important to note that for classification purposes, stimulus presentation was not jittered and alpha power is modulated by pre-stimulus preparatory attention (e.g.^51^). We have similarly found evidence for putative preparatory encoding state engagement when trial times are fixed^34^ as compared to variable^12^. Thus, together with univariate effects of stimulus predictability on attentional engagement, these findings suggest that individuals may modulate brain state engagement on sub-second timescales in response to top-down and bottom-up inputs.

Across these three tasks, we show that an independently validated, cross-study mnemonic state classifier can detect moment to moment shifts in attention. Given the absence of long-term episodic memory demands in these tasks, our interpretation is that memory encoding and memory retrieval brain states reflect external and internal attention respectively. Most importantly, we reveal that mnemonic state engagement can change in a matter of milliseconds. These fluctuations have critical implications for growing research on how the brain switches between external and internal attention states^52–54^. Although in the course of laboratory research we typically divide trials into conditions of interest, this is not how brain states are likely to be engaged outside the lab. In everyday life we must frequently shift between internal attention – thinking about what you’ll have for dinner – and external attention – the oncoming car approaching you at the intersection. The need to switch between mnemonic brain states impacts how they are recruited^55, 56^ and it may be more difficult to switch from an external to internal state^52^. Thus, mnemonic state classification of electrophysiological signals provides a critical avenue for assessing moment-to-moment changes in attention.

The relationship between moment-to-moment brain state engagement and behavior will ultimately depend on both top-down task goals and bottom-up stimulus factors. For instance, relying on prior knowledge to elaborate a representation (e.g. depth of processing,^57^) is an instance in which internal attention could facilitate memory formation through accessing of stored semantic knowledge. However, in other cases, recruiting the retrieval state can impair subsequent memory for to-be-encoded items^9^. In healthy aging, over-reliance on prior knowledge may come at the expense of attention to external, episodic details^58^ and healthy older adults may be biased to engage the retrieval state even when their goal is to encode^59, 60^. Thus it will be critical to determine how the precise temporal dynamics of brain state engagement supports cognitive processes and behavior.

In conclusion, mnemonic brain states are robustly modulated by external and internal attention demands across a variety of task factors. Thus mnemonic brain states reflect domain general, content-independent attentional states. We can therefore use online, realtime measurements of mnemonic states to detect, select and correct attentional state engagement.

## Online Methods

### Data and Code Availability

The raw, de-identified data and the associated experimental and analysis codes used in this study will be made available via the Long Term Memory laboratory website (https://longtermmemorylab.com) upon publication. All analyses were pre-registered (osf.io/x7jvs) unless noted otherwise.

### Participants

48 adults (30 female, age range = 18-31, mean age = 20 years) fluent English speakers from the University of Virginia (UVA) community participated. Participants were recruited from flyers posted around the UVA campus and surrounding areas. Our sample size was determined *a priori* as described in the pre-registration report of this study (osf.io/x7jvs). All participants had normal or corrected-to-normal vision. Informed consent was obtained in accordance with the UVA Institutional Review Board for Social and Behavioral Research and participants were compensated for their participation. Four participants were excluded from the final dataset: one due to poor signal quality, one due to failure to comply with task instructions, one due to data loss resulting from a power outage, and one whose mean accuracy on the WMT exceeded the lower threshold based on the group data (Mean (2.5*SD)). We report results from the remaining 44 participants.

### Materials

*Overview.* Each participant in this experiment completed three distinct tasks, each designed to elicit internal and external attention (Figure 1). Task order was counterbalanced across participants. Participants completed a brief practice phase before beginning each task.

### Visual Attentional Task (VAT)

The design of this task parallels the mnemonic state task that we have previously employed to detect memory encoding and memory retrieval states^9–12^. The stimuli and timing parameters are the same; the specific task cues that participants encountered were modified to direct participants to selectively engage in either external or internal attention.

#### Stimuli

Stimuli consisted of images of common objects drawn from an image database with multiple exemplars per object category^61^. Participants encountered a total of 192 exemplars from 192 unique object categories. We created duplicate images of all objects and converted the duplicates to grayscale. We removed potentially ambiguous images which tended to be those that are entirely metal (e.g. a strainer). Object condition assignment was randomly generated for each participant.

#### Experimental Design

In each of eight runs, participants viewed 24 object images. Half of the objects were presented in grayscale and half were presented in color (Figure 1A). On each trial, participants saw a cue word, either “LOOK” or “IMAGINE” for 500 ms. The cue was followed by a 1500 ms delay interval, followed by presentation of an object for 2000 ms. Each trial was separated by a 1000 ms inter-trial interval (ITI). On trials with a “LOOK” instruction (external trials), participants were to look at the presented object. On trials with an “IMAGINE” instruction (internal trials), participants were to try to imagine the presented object as the opposite – color or grayscale – of how it was presented. That is, if the presented object was a grayscale bench, in the external condition participants should focus on the grayscale bench vs. in the internal condition they should imagine the grayscale bench in color. Participants did not make any behavioral responses.

Across the eight runs, participants encountered a total of 192 trials, 96 LOOK/external and 96 IMAGINE/internal. Cue condition was random across trials.

### Working Memory Task (WMT)

This task is a modified version of a previously reported pre- and retro-cued working memory task^49^.

#### Stimuli

Stimuli consisted of X’s, unfilled boxes, and an asterisk. A central fixation cross was present throughout all trial intervals. X’s could be one of four colors: red, yellow, green, or blue. Four X’s presented during the array interval were located at one of four locations (upper left, upper right, lower left, lower right) relative to the fixation cross. A single X was presented centrally during the probe interval. Unfilled boxes served as the pre and retro cues. Two shaded lines were overlaid on the unfilled boxes to cue one of the four locations.

#### Experimental Design

In each of six runs, participants completed 32 trials. Each trial had five intervals (Figure 1B). During the trial onset interval, an asterisk was presented for 200 ms. During the pre-cue interval, an unfilled box was presented for 100 ms. During the array interval, four X’s were presented for 500 ms. During the retro-cue interval, an unfilled box was presented for 100 ms. During the probe interval, a single X was presented for 100 ms. Each interval was separated by a fixed inter-stimulus interval (ISI) the duration of which was calibrated to make each interval (stimulus presentation plus fixation) total 2000 ms. Thus, the trial onset interval was followed by an 1800 ms ISI, the pre and retro cue intervals were followed by 1900 ms ISIs, the array interval was followed by a 1500 ms ISI, and the probe interval was followed by a 1900 ms ISI. Participant responses were accepted during the final 1900 ms ISI following the probe interval.

We manipulated *cue type*: pre-cued vs. retro-cued; *cue validity* : valid, invalid; *probe color* : red, yellow, green, blue; and *cue location*: upper left, upper right, lower right, lower left. Within each run, half of the trials were pre-cued (n=16) and half of the trials were retro-cued (n=16). For pre-cued trials, the cue appeared during the pre-cue interval between trial onset and the array interval. For retro-cued trials, the cue appeared during the retro-cue interval between the array and probe intervals. The other interval (retro-cue or pre-cue, respectively) had an unfilled square presented to equate visual presentation across the two intervals. The pre- and retro-cued trials were further subdivided into validly cued (n=8) and invalidly cued (n=8). Finally, the trials were subdivided again to ensure an equal number of cued and probed colors within each cue type and cue validity condition (n=2 for each of red, yellow, green, blue). Cue locations were not strictly balanced as a full balance would require 64 trials per run which in our opinion yields runs that are too long (approximately five minutes) for participants to maintain adequate focus. Instead, we ensured that there were equal numbers of cued locations (n=2 of each) within each cue validity condition.

Participants’ task was to respond if the probed color had appeared in the (pre or retro) cued location. Thus the correct responses were “yes,” for valid trials and “no,” for invalid trials. Participants responded via key press and response options were counterbalanced across participants. Participants were instructed to respond as quickly and accurately as possible and were given a maximum of 1900 ms to respond before the experiment proceeded to the next trial.

Across the six runs, participants encountered a total of 192 trials, 96 pre-cue/external and 96 retro- cue/internal. Cue type was random across trials.

#### Rationale for Design Alterations

We modified several parameters of the original task^49^. First, we shortened and fixed the ISI between the components of each trial. We shortened the ISIs as EEG is not constrained by the slowness of hemodynamic response and shorter ISIs allow a greater number of trials to be collected from each participant. We used fixed rather than jittered ISIs to facilitate both multivariate decoding and cross-trial and cross-task comparisons. Fixed ISIs do allow participants to anticipate upcoming stimuli; however, we did not expect this to selectively affect any particular condition. Therefore, we did not anticipate that fixed ISIs would impact our expected outcomes. Second, we increased the duration of the array from 100 ms to 500 ms. This longer duration matches a working memory study from our lab^35^ and provides a longer interval to assess stimulus-related external attention. Third, we increased the number of trials per run (from 24 to 32) and total runs (from 4 to 6). We increased trial and run counts to both facilitate condition balancing and to equate the number of internal/external trials across tasks.

### Auditory Attentional Task (AAT)

This task is a modified version of previously reported externally vs. internally directed attention tasks^24, 25^.

#### Stimuli

Stimuli were standard tones (800 Hz) and target tones (1000 Hz) played for 200 ms. Within each block, standard tones occurred on 80% of trials and target tones occurred on 20% of trials.

#### Experimental Design

In each of four runs, participants completed two blocks of 24 trials each (Figure 1C). Each block began with a get ready screen (2000 ms), followed by an instruction screen (2000 ms), and an ISI (1000 ms). Each trial began with a tone played for 200 ms. Trials were separated by a fixed ITI of 1800 ms during which responses were accepted. In the *Externally Directed Attention* (EDA) blocks, participants were instructed to focus on the tones and respond via key press when they heard a target tone. In the *Internally Directed Attention* (IDA) blocks, participants were instructed to focus on their thoughts and provided no responses. Block order was counterbalanced across runs. Tone presentation was random except that target tones were prevented from being presented on the first trial of any block. Participants were encouraged to respond as quickly and accurately as possible within the ITI.

Across the four runs, participants encountered a total of 192 trials, 96 EDA and 96 IDA. Task condition was blocked within run.

#### Rationale for Design Alterations

We modified several parameters of the original task^24, 25^. We reduced the number of trials per run (from 25 per condition to 24 per condition) and the number of runs (from a maximum of 28 to 4). We reduced trial and run counts to match the number of internal/external trials across tasks. As with the Working Memory Task, we used a fixed, rather than jittered, ITI, to facilitate multivariate decoding and cross-trial, cross-task comparisons. We increased the ITI to match the other tasks.

### EEG Data Acquisition and Preprocessing

EEG recordings were collected using a BrainVision system and an ActiCap equipped with 64 Ag/AgCl active electrodes positioned according to the extended 10-20 system. All electrodes were digitized at a sampling rate of 1000 Hz and were referenced to electrode FCz. Offline, electrodes were later converted to an average reference. Impedances of all electrodes were kept below 50kΩ. Electrodes that demonstrated high impedance or poor contact with the scalp were excluded from the average reference; however, these electrodes were included in all subsequent analysis steps. Bad electrodes were determined by voltage thresholding (see below).

Custom python codes were used to process the EEG data. We applied a high pass filter at 0.1 Hz, followed by a notch filter at 60 Hz and harmonics of 60 Hz to each participant’s raw EEG data. We then performed three preprocessing steps^62^ to identify electrodes with severe artifacts. First, we calculated the mean correlation between each electrode and all other electrodes as electrodes should be moderately correlated with other electrodes due to volume conduction. We z-scored these means across electrodes and rejected electrodes with z-scores less than -3. Second, we calculated the variance for each electrode, as electrodes with very high or low variance across a session are likely dominated by noise or have poor contact with the scalp. We then z-scored variance across electrodes and rejected electrodes with a |z*| >* = 3. Finally, we expect many electrical signals to be autocorrelated, but signals generated by the brain versus noise are likely to have different forms of autocorrelation. Therefore, we calculated the Hurst exponent, a measure of long-range autocorrelation, for each electrode and rejected electrodes with a |z*| >* = 3. Electrodes marked as bad by this procedure were excluded from the average re-reference. We then calculated the average voltage across all remaining electrodes at each time sample and re-referenced the data by subtracting the average voltage from the filtered EEG data. We used wavelet-enhanced independent component analysis^63^ to remove artifacts from eyeblinks and saccades.

### EEG Data Analysis

We applied the Morlet wavelet transform (wave number 6) to the entire EEG time series across electrodes, for each of 46 logarithmically spaced frequencies (2-100 Hz;^11^), across all experiments. After log-transforming the power, we downsampled the data by taking a moving average across 100 ms time intervals from either 500 ms preceding to 2000 ms following stimulus onset for the VAT and AAT or from 2000 ms preceding to 4000 ms following stimulus onset for the WMT. We used a sliding window of 25 ms, resulting in 97 time intervals (25 non-overlapping) for the VAT and AAT and 317 time intervals (80 non-overlapping) for the WMT. Mean and standard deviation power were calculated across all trials and across time points for each frequency. Power values were then z-transformed by subtracting the mean and dividing by the standard deviation power.

### Univariate Analyses

We performed three exploratory univariate contrasts. We compared spectral signals averaged over the stimulus interval for external (LOOK, EDA) vs. internal (IMAGINE, IDA) trials in the VAT and AAT. Additionally, we compared spectral signals averaged over the pre-cue interval for external (pre-cued) trials vs. spectral signals averaged over the retro-cue interval for internal (retro-cued) trials in the WMT. For each contrast, participant, electrode, and frequency, we calculated zPower in each of the two conditions and averaged zPower over the 2000 ms interval of interest. For each task (VAT, WMT, AAT) and each participant, we generated a “template” spectral patterns across 63 electrodes *×* 46 frequencies. We then extracted trial level patterns in the other two tasks. We used Pearson correlations to relate trial level external or internal patterns with the external or internal “template” pattern. All rho values were Fisher-Z transformed and averaged across trials. This analysis yielded a single average correlation value for each participant and pair of tasks.

### Pattern Classification Analyses

Pattern classification analyses were performed using penalized (L2) logistic regression implemented via the sklearn module in Python and custom Python code. For all classification analyses, classifier features were comprised of spectral power across 63 electrodes and 46 frequencies. Before pattern classification analyses were performed, an additional round of z-scoring was performed across features (electrodes and frequencies) to eliminate trial-level differences in spectral power^11^. Therefore, mean univariate activity was matched precisely across all conditions and trial types. Our classification approach yields two outputs, “classification accuracy,” and “classification evidence.” “Classification accuracy” represents a binary coding of whether the classifier successfully guessed the attention label, external or internal. We used classification accuracy to assess both within-task and cross-task classifier performance (i.e. whether external/internal trials could be decoded). Classification evidence is a continuous value reflecting the logit-transformed probability that the classifier assigns a specific label (e.g. encode, retrieve) to each trial. We used classification evidence as a trial-specific, continuous measure of mnemonic state information.

#### Within-task classification analysis

To assess the extent to which internal and external attentional states are discriminable, we conducted leave-one-run-out cross-validated-classification (penalty parameter = 1) separately for each of the three tasks. We trained each within-participant classifier to distinguish external and internal trials based on z-scored spectral power averaged over the stimulus interval (2000 ms) in the VAT and AAT and z-scored spectral power from the pre-cue interval of pre-cued trials and retro-cue interval of retro-cued trials (2000 ms) in the WMT. The rationale for selecting these specific WMT trials is that the visual stimulus input is the same across the two trials: both have a cue presented for 100 ms followed by a 1900 ms delay. Thus, decoding success is more attributable to process differences between pre-cued and retro-cued trials given the matched visual input.

#### Cross-task classification analysis

To test the extent to which external and internal states generalize across stimulus modality and forms of attention, we conducted cross-task classification (penalty parameter = 1) on the stimulus interval data from the VAT and AAT and on the pre/retro cue interval data from the WMT. We ran six sets of within-participant classifiers to fully cross training and test data. Classifiers were trained to distinguish external and internal trials based on z-scored spectral power from either the stimulus interval of the VAT or AAT or the pre and retro-cue intervals from the pre/retro-cued trials in the WMT. Classifiers were tested on data from one of the other two tasks. We averaged classification accuracy values across each pair of tasks, yielding classification accuracy values for VAT-AAT, VAT-WMT, and AAT-WMT.

#### Cross study mnemonic state classification

To measure mnemonic state engagement across the three tasks, we conducted three stages of classification using the same methods as in our prior work^6, 12^. First, we conducted within participant leaveone-run-out cross-validated classification (penalty parameter = 1) on all participants who completed the mnemonic state task (N = 143, see refs.^36^ and^12^ for details). The classifier was trained to distinguish encoding vs. retrieval states based on spectral power averaged across the 2000 ms stimulus interval during List 2 trials. For each participant, we generated true and null classification accuracy values. We permuted condition labels (encode, retrieve) for 1000 iterations to generate a null distribution for each participant. Any participant whose true classification accuracy fell above the 90th percentile of their respective null distribution was selected for further analysis (N = 57). Second, we conducted leave-one-participant-out cross-validated classification (penalty parameter = 0.0001) on the selected participants to validate the mnemonic state classifier and obtained classification accuracy of 59.29% which is significantly above chance (*t* _56_ = 7.667, p *<* 0.0001), indicating that the cross-subject mnemonic state classifier is able to distinguish encoding and retrieval states. Finally, we applied the cross-subject mnemonic state classifier to all trials within the current study. This approach provides a trial-level estimate of mnemonic state evidence across the three tasks.

#### Statistical Analyses

To assess attentional state decoding accuracy, we used paired-sample *t* -tests to compare within-task classification accuracy across participants to chance-level decoding accuracy, as determined by permutation procedures. Namely, for each task and each participant, we shuffled the condition labels of interest (e.g., external and internal) and then calculated classification accuracy. We repeated this procedure 1000 times for each participant for each task and then averaged the 1000 shuffled accuracy values for each participant. These mean values were used as participant-specific empirically derived measures of chance accuracy. We utilized the same general procedure to assess cross-task generalization. We used repeated measures ANOVAs (rmANOVAs) and *t* -tests to assess the effect of condition (external, internal) on mnemonic state evidence over time.

We assessed significance of the univariate contrasts through cluster-based correction^64^. For each of the three contrasts, we performed paired samples *t* -tests at each electrode *×* frequency point across participants. True *p* values that exceeded an uncorrected threshold of *p <* 0.05 were divided into contiguous clusters, separately for positively vs. negatively signed *t* values. To obtain chance values, we permuted the participant-level average zPower values across conditions and generated associated chance *t* and *p* values. As with the true *p* values, any chance values within a given permutation iteration (1000 total) that exceeded the uncorrected threshold were divided into contiguous clusters. *T* -values were summed within each cluster and the maximum test statistic was retained for the iteration. Maximum *t* -values were aggregated into a single distribution and true clusters with *t* -values that exceeded the 97.5th percentile (accounting for the separate testing of positive and negative clusters) were marked as significant.

We used one sample *t* -tests to compare template zRho correlations to zero.

We applied multiple comparisons corrections for post-hoc tests (using False Discovery Rate correction;^65^). We conducted Bayes factor analysis using the BayesFactor package (version 0.9.12-4.8) in R (version 4.4.0) with the default prior settings. We used the function ttestBF to compare models.

## Acknowledgments

This work was supported by a grant from the National Institutes of Health (NINDS R01 NS132872, PI: N.M.L.). We thank Eddie Ashie, Matthew Bair, Hannah Buras, Leah Cassidy, Lola Fuentes Brock, Rithik Kalikota, Amanda Smith, Salonee Verma, and Theo Wilson for assistance with data collection.

## Notes

Conflict of Interest: The authors declare no competing financial interests.

### Competing Interest Statement

The authors have declared no competing interest.

## References

1 Chun, M. M. and Turk-Browne, N. B. (2007). Interactions between attention and memory. Current Opinion in Neurobiology 17(2), 177–184.

2 Nobre, A. C. Attention. In Steven’s Handbook of Experimental Psychology and Cognitive Neuroscience, Wixted, J. T., editor. John Wiley & Sons, Inc. fourth edition (2018).

3 Chun, M. M., Golomb, J. D., and Turk-Browne, N. B. (2011). A taxonomy of external and internal attention. Annual Review of Psychology 62, 73–101.

4 Verschooren, S. and Egner, T. (2023). When the mind’s eye prevails: The internal dominance over external attention (idea) hypothesis. Psychonomic Bulletin & Review 30, 1668–1688.

5 Logan, G. D., Lilburn, S. D., Ulrich, J. E., Weeks, E. E., and Koo, R. (2026). Spotlight on the past: Focusing attention on long-term memory. Memory & Cognition .

6 Long, N. M. (2023). The intersection of the retrieval state and internal attention. Nature Communications 14(3861).

7 Cleary, A. M., McNeely-White, K. L., Neisser, J., Drane, D. L., Liégeois-Chauvel, C., and Pedersen, N. P. (2025). Does familiarity-detection flip attention inward? the familiarity-flip-of-attention account of the primacy effect in memory for repetitions. Memory & Cognition 53.

8 Hasselmo, M. E., Bodelón, C., and Wyble, B. P. (2002). A proposed function for hippocampal theta rhythm: Separate phases of encoding and retrieval enhance reversal of prior learning. Neural Computation 14(4), 793–817.

9 Long, N. M. and Kuhl, B. A. (2019). Decoding the tradeoff between encoding and retrieval to predict memory for overlapping events. NeuroImage 201.

10 Long, N. M. and Kuhl, B. A. (2021). Cortical representations of visual stimuli shift locations with changes in memory states. Current Biology 31(5).

11 Smith, D. E., Moore, I. L., and Long, N. M. (2022). Temporal context modulates encoding and retrieval of overlapping events. Journal of Neuroscience 42(14), 3000–3010.

12 Han, S. and Long, N. M. (2025). Evidence for a reactionary account of retrieval state initiation. Imaging Neuroscience 3.

13 Poulet, J. F. A. and Petersen, C. C. H. (2008). Internal brain state regulates membrane potential synchrony in barrel cortex of behaving mice. Nature 454(7206), 881–885.

14 Harris, K. D. and Thiele, A. (2011). Cortical state and attention. Nature Reviews Neuroscience 12(9), 509–523.

15 Kay, K. and Frank, L. M. (2019). Three brain states in the hippocampus and cortex. Hippocampus 29, 184–238.

16 Corbetta, M. and Shulman, G. L. (2002). Control of goal-directed and stimulus-driven attention in the brain. Nature Reviews Neuroscience 3(3), 201–215.

17 Yeo, B. T., Krienen, F. M., Sepulcre, J., Sabuncu, M. R., Lashkari, D., Hollinshead, M., Roffman, J. L., Smoller, J. W., Zö llei, L., Polimeni, J. R., et al. (2011). The organization of the human cerebral cortex estimated by intrinsic functional connectivity. Journal of Neurophysiology 106(3), 1125–1165.

18 Kim, H. (2014). Involvement of the dorsal and ventral attention networks in oddball stimulus processing: A meta-analysis. Human Brain Mapping 35(5), 2265–2284.

19 Müller, M. M., Gruber, T., and Keil, A. (2000). Modulation of induced gamma band activity in the human eeg by attention and visual information processing. International Journal of Psychophysiology 38, 283–300.

20 Helfrich, R. F., Breska, A., and Knight, R. T. (2019). Neural entrainment and network resonance in support of top-down guided attention. Current Opinion in Psychology 29, 82–89.

21 Raichle, M. E., MacLeod, A. M., Snyder, A. Z., Powers, W. J., Gusnard, D. A., and Shulman, G. L. (2001). A default mode of brain function. Proceedings of the National Academy of Sciences of the United States of America 98(2), 676–682.

22 Buckner, R. L. and DiNicola, L. M. (2019). The brain’s default network: updated anatomy, physiology and evolving insights. Nature Reviews Neuroscience 20, 593–608.

23 Cooper, N. R., Croft, R. J., Dominey, S. J. J., Burgess, A. P., and Gruzelier, J. H. (2003). Paradox lost? Exploring the role of alpha oscillations during externally vs. internally directed attention and the implications for idling and inhibition hypotheses. International Journal of Psychophysiology 47(1), 65– 74.

24 Kam, J. W. Y., Solbakk, A.-K., Endestad, T., Meling, T. R., and Knight, R. T. (2018). Lateral prefrontal cortex lesion impairs regulation of internally and externally directed attention. NeuroImage 175, 91–99.

25 Kam, J. W. Y., Lin, J. J., Solbakk, A.-K., Endestad, T., Larsson, P. G., and Knight, R. T. (2019). Default network and frontoparietal control network theta connectivity supports internal attention. Nature Human Behavior 3, 1263–1270.

26 Johnson, J. A. and Zatorre, R. J. (2005). Attention to simultaneous unrelated auditory and visual events: Behavioral and neural correlates. Cerebral Cortex 15(10), 1609–1620.

27 Mazaheri, A., van Schouwenburg, M. R., Dimitrijevic, A., Denys, D., Cools, R., and Jensen, O. (2014). Region-specific modulations in oscillatory alpha activity serve to facilitate processing in the visual and auditory modalities. NeuroImage 87, 356–362.

28 Dixon, M. L., Fox, K. C., and Christoff, K. (2014). A framework for understanding the relationship between externally and internally directed cognition. Neuropsychologia 62, 321–330.

29 Dijkstra, N., Bosch, S. E., and van Gerven, M. A. J. (2019). Shared neural mechanisms of visual perception and imagery. Trends in Cognitive Sciences 23(5).

30 Tulving, E. *Elements of Episodic Memory*. Oxford, New York, (1983).

31 Tarder-Stoll, H., Jayakumar, M., Dimsdale-Zucker, H. R., Günseli, E., and Aly, M. (2020). Dynamic internal states shape memory retrieval. Neuropsychologia 138.

32 Mulligan, N. W., Spataro, P., and West, J. T. (2023). Memory and attention: A double dissociation between memory encoding and memory retrieval. Cognition 238.

33 Dede, A. J., Cross, Z. R., Gray, S. M., Kelly, J. P., Yin, Q., Vahidi, P., Asano, E., Schuele, S. U., Rosenow, J. M., Wu, J. Y., et al. (2025). Episodic memory involves transient and sparse connectivity aligned to both internal and external events. PLoS Biology 23(11).

34 Bair, M. B. and Long, N. M. (2026). A shared brain state for episodic and semantic retrieval. bioRxiv.

35 Nguyen, D. T. and Long, N. M. (2026). Working memory demands modulate memory brain state engagement. bioRxiv .

36 Hong, Y., Smith, D. E., Moore, I. L., and Long, N. M. (2023). Spatiotemporal dynamics of memory encoding and memory retrieval states. Journal of Cognitive Neuroscience 35(9).

37 Fiebelkorn, I. C., Saalmann, Y. B., and Kastner, S. (2013). Rhythmic sampling within and between objects despite sustained attention at a cued location. Current Biology 23, 2553–2558.

38 Wiesman, A. I. and Wilson, T. W. (2020). Attention modulates the gating of primary somatosensory oscillations. NeuroImage 211.

39 Galas, L., Donovan, I., and Dugué, L. (2025). Attention rhythmically shapes sensory tuning. Journal of Neuroscience 45(7).

40 Schupp, H. T., Lutzenberger, W., Birbaumer, N., Miltner, W., and Braun, C. (1994). Neurophysiological differences between perception and imagery. Cognitive Brain Research 2, 77–86.

41 Villena-González, M., López, V., and Rodríguez, E. (2016). Orienting attention to visual or verbal/auditory imagery differentially impairs the processing of visual stimuli. NeuroImage 132, 71–78.

42 Baddeley, A. D. and Hitch, G. J. Working memory. In The psychology of learning and motivation: Advances in research and theory, Bower, G. H., editor, volume 8, 47–90. Academic Press, New York (1974).

43 Desimone, R. and Duncan, J. (1995). Neural mechanisms of selective visual attention. Annual Review of Neuroscience 18, 193–222.

44 Jensen, O., Bonnefond, M., and VanRullen, R. (2012). An oscillatory mechanism for prioritizing salient unattended stimuli. Trends in Cognitive Sciences 16(4), 200–206.

45 Samaha, J., Sprague, T. C., and Postle, B. R. (2016). Decoding and reconstructing the focus of spatial attention from the topography of alpha-band oscillations. Journal of Cognitive Neuroscience 28(8), 1090–1097.

46 Bonnefond, M. and Jensen, O. (2013). Alpha oscillations serve to protect working memory maintenance against anticipated distracters. Current Biology 22(20).

47 Lavie, N., Hirst, A., Viding, E., and de Fockert, J. W. (2004). Load theory of selective attention and cognitive control. Journal of Experimental Psychology: General 133(3), 339–354.

48 Pillay, S., Durgerian, S., and Sabri, M. (2016). Perceptual demand and distraction interactions mediated by task-control networks. NeuroImage 138.

49 Nobre, A. C., Coull, J. T., Maquet, P., Frith, C. D., Vandenberghe, R., and Mesulam, M. M. (2004). Orienting attention to locations in perceptual versus mental representations. Journal of Cognitive Neuroscience 16(3), 363–373.

50 Myers, N. E., Stokes, M. G., and Nobre, A. C. (2017). Prioritizing information during working memory: Beyond sustained internal attention. Trends in Cognitive Sciences 21(6).

51 Hanslmayr, S., Aslan, A., Staudigl, T., Klimesch, W., Herrmann, C. S., and Bä uml, K.-H. (2007). Prestimulus oscillations predict visual perception performance between and within subjects. NeuroImage 37, 1465–1473.

52 Verschooren, S., Pourtois, G., and Egner, T. (2020). More efficient shielding for internal than external attention? evidence from asymmetrical switch costs. Journal of Experimental Psychology: Human Perception and Performance 46(9), 912–925.

53 Poskanzer, C. and Aly, M. (2023). Switching between external and internal attention in hippocampal networks. Journal of Neuroscience 43(38), 6538–6552.

54 Nobre, A. C. and Gresch, D. (2025). How the brain shifts between external and internal attention. Neuron 113.

55 Morcom, A. M. and Rugg, M. D. (2002). Getting ready to remember: the neural correlates of task set during recognition memory. NeuroReport 13(1).

56 Evans, L. H., Williams, A. N., and Wilding, E. L. (2015). Electrophysiological evidence for retrieval mode immediately after a task switch. NeuroImage 108, 435–440.

57 Craik, F. I. M. and Tulving, E. (1975). Depth of processing and the retention of words in episodic memory. Journal of Experimental Psychology: General 104(3), 268–294.

58 Lalla, A., Tarder-Stoll, H., Hasher, L., and Duncan, K. D. (2022). Aging shifts the relative contributions of episodic and semantic memory to decision-making. Psychology and Aging 37(6), 667–680.

59 Moore, I. L., Smith, D. E., and Long, N. M. (2025). Mnemonic brain state engagement is diminished in healthy aging. Neurobiology of Aging 151.

60 Buras, H. R., Han, S., and Long, N. M. (2026). Healthy aging, processing speed, and mnemonic brain state engagement. bioRxiv .

61 Konkle, T., Brady, T. F., Alvarez, G. A., and Oliva, A. (2010). Conceptual distinctiveness supports detailed visual long-term memory for real-world objects. Journal of Experimental Psychology: General 139(3), 558–578.

62 Nolan, H., Whelan, R., and Reilly, R. (2010). Faster: fully automated statistical thresholding for eeg artifact rejection. Journal of neuroscience methods 192(1), 152–162.

63 Castellanos, N. P. and Makarov, V. A. (2006). Recovering EEG brain signals: Artifact suppression with wavelet enhanced independent component analysis. Journal of Neuroscience Methods 158(2), 300–312.

64 Maris, E. and Oostenveld, R. (2007). Nonparametric statistical testing of EEG- and MEG-data. Journal of Neuroscience Methods 164, 177–190.

65 Benjamini, Y. and Hochberg, Y. (1995). Controlling the false discovery rate: a practical and powerful approach to multiple testing. Journal of the Royal Statistical Society. Series B (Methodological) 57(1), 289–300.

